# An X Chromosome Transcriptome Wide Association Study Implicates ARMCX6 in Alzheimer’s Disease

**DOI:** 10.1101/2023.06.06.543877

**Authors:** Xueyi Zhang, Lissette Gomez, Jennifer Below, Adam Naj, Eden Martin, Brian Kunkle, William S. Bush

## Abstract

**Background:** The X chromosome is often omitted in disease association studies despite containing thousands of genes which may provide insight into well-known sex differences in the risk of Alzheimer’s Disease.

**Objective:** To model the expression of X chromosome genes and evaluate their impact on Alzheimer’s Disease risk in a sex-stratified manner.

**Methods:** Using elastic net, we evaluated multiple modeling strategies in a set of 175 whole blood samples and 126 brain cortex samples, with whole genome sequencing and RNA-seq data. SNPs (MAF>0.05) within the *cis*-regulatory window were used to train tissue-specific models of each gene. We apply the best models in both tissues to sex-stratified summary statistics from a meta-analysis of Alzheimer’s disease Genetics Consortium (ADGC) studies to identify AD-related genes on the X chromosome.

**Results:** Across different model parameters, sample sex, and tissue types, we modeled the expression of 217 genes (95 genes in blood and 135 genes in brain cortex). The average model R^2^ was 0.12 (range from 0.03 to 0.34). We also compared sex-stratified and sex-combined models on the X chromosome. We further investigated genes that escaped X chromosome inactivation (XCI) to determine if their genetic regulation patterns were distinct. We found ten genes associated with AD at p < 0.05, with only *ARMCX6* in female brain cortex (p = 0.008) nearing the significance threshold after adjusting for multiple testing (α = 0.002).

**Conclusions:** We optimized the expression prediction of X chromosome genes, applied these models to sex-stratified AD GWAS summary statistics, and identified one putative AD risk gene, *ARMCX6*.

## Introduction

Nearly two-thirds of individuals with Alzheimer’s Disease (AD) are women [1], as many other human traits also present sex disparities. While these differences may be due to systematic exposure to sex hormones or other endogenous factors, X chromosome genetic variation and its expression are potential contributing factors. The human X chromosome is composed of 155 million base pairs and contains thousands of genes. The copy number of the X chromosome defines biological sex. While females have two copies, genes on one X chromosome are randomly silenced via DNA methylation in a process called X-chromosome inactivation (XCI). The XCI patterns vary from person to person, with a proportion of genes escaping the silencing process [2–5]. The choice of the silenced copy within a cell is random early in embryo development. At the same time, X-linked genes’ mosaic expression at the individual level can be skewed towards the maternal or paternal copy of the gene [6,7]. Functionally, X-linked genes are responsible for various processes, most notably human cognition and the development of multiple tissues, including neural and bone [8].

Despite its potential importance, X chromosome variants are typically removed from genome-wide association studies (GWAS) during quality control due to the complexity of accounting for differences in chromosomal copy number between males and females. In general, methods used for autosomal genotypes cannot be applied directly to X chromosome variants as they do not account for the genetic imbalance or expected X-inactivation. [9–11] As a result, only one-third of genome-wide association studies include evaluations of variants on the X [12]. None of the recent large sample AD GWAS [13–17] included X chromosome. However, one smaller MRI-based study (N = 931) by Homann et al. [18] is notable as they meta-analyzed sex-specific X chromosome associations. The number of reported GWAS associations on the X chromosome is smaller than that of autosomes [19], and Kukurba et al. discovered a relative depletion of *cis*-eQTLs on the X chromosome using a general eQTL-mapping pipeline on both autosomes and the X chromosome [20]. These differences can either result from the inefficient power of the tests or the underlying different biological mechanisms and genetic architecture between autosomes and the X chromosome. The exclusion of X chromosome variants also propagates into the post-GWAS era. Expression QTL (quantitative trait loci) and other studies of molecular phenotypes also often exclude the sex chromosomes. MetaBrain [21], which harmonized RNA-seq samples from 14 studies and prioritized *cis-* and *trans-*eQTLs for brain-related traits, also did not report any loci on the X chromosome. Similarly, the largest harmonized blood *cis-* and *trans-* eQTLs resource, eQTLGEN [22], also excluded the X chromosome. Popular extensions of GWAS, such as transcriptome-wide association studies (TWAS) [23,24], omit the X chromosome due to lack of adequate models of X-linked gene expression. Tissue-specific and cross-tissue TWAS on AD [25,26] did not include the X chromosome.

In this study, we evaluated multiple approaches to fit expression prediction models for X chromosome genes and produced a set of sex-stratified X-chromosome prediction models. By applying these models in a TWAS on AD GWAS summary statistics, we detected one gene, *ARMCX6*, borderline associated with AD risk in the cortex in female-only samples.

## Methods

### Data resources

We retrieved WGS and RNA-seq expression levels data (phs000424.v7.p2) from the brain cortex and whole blood sample from the GTEx data [27] to train the prediction models. We selected cortex because of the availability and larger sample size of the validation dataset, in addition to its direct relevance to the AD disease process. Whole blood is the most accessible tissue for transcriptome evaluations and also captures immune cell expression, which is also likely relevant for AD. For validation, we used (1) whole-genome imputed genotype data and whole blood sample RNA-seq data from the DGN (Depression susceptibility Genes and Network) expression cohort [28] and (2) WGS and temporal cortex (TCX) RNA-seq data from the MayoRNAseq study [29]. Gene expression data of the MayoRNAseq cohort has already been adjusted with Alzheimer’s Disease (AD) diagnosis, sex, and age at death.

The GTEx donors were from post-mortem autopsy or organ and tissue transplantation settings. The DGN cohort reported no differences in eQTL detection between depression cases and controls. [28] MayoRNAseq study included patients with AD and related dementias and health control on donors from the Mayo Clinic Brain Bank and Banner Health. [29] These datasets can represent typical patterns of gene expression in blood and cortex.

The key features of the three study populations are listed in Table 1. DGN and MayoRNAseq only recruited subjects of European descent from the United States. To evaluate different modeling strategies in a homogeneous study population and make generalizable recommendations, we restricted the samples to the GTEx participants who were self-identified as non-Hispanic white (NHW). The age data in GTEx and MayoRNAseq is the age at death of the donors, while that for DGN is the age at sampling. MayoRNAseq subjects over 90 years old (recorded as 90+) were treated as age 91. It is noteworthy that the three studies exhibit different sex ratios, with males comprising two-thirds of the GTEx NHW cohort, females comprising two-thirds of the DGN cohort, and the MayoRNAseq TCX cohort displaying gender balance.

**Table 1.** Overview of the study datasets for training and validating the models.

|  | GTEx blood | GTEx cortex | DGN blood | MayoRNAseq cortex |
| --- | --- | --- | --- | --- |
| Sample size | 175 | 126 | 917 | 255 |
| Female | 58 | 46 | 648 | 130 |
| Male | 117 | 80 | 269 | 125 |
| Age | 56.1(10.5) | 57.3(10.6) | 44.7(10.9) | 80.5(8.5) |
| Female | 57.4(9.0) | 57.9(11.4) | 45.0(10.5) | 81.5(8.5) |
| Male | 55.5(11.2) | 57.0(10.3) | 44.1(11.7) | 79.4(8.3) |
| Platform | WGS<br>RNA-seq | WGS<br>RNA-seq | Genotype<br>RNA-seq | WGS<br>RNA-seq |
| Genome Assembly | GRCh37 | GRCh37 | GRCh37 | GRCh37 |
| Number of expressed genes | 600 | 766 | 486 | 587 |

Additionally, MayoRNAseq subjects are significantly older than GTEx and DGN. Human subjects and ethics statements for these studies are available in their respective publications [27–29].

### Data processing

To keep the analysis concise, we dropped pseudoautosomal regions (PARs) (GRCh37, X:60,001-2,699,520, and X:154,931,044-155,260,560) for this study. The PAR boundaries were retrieved from https://www.ncbi.nlm.nih.gov/grc/human. Genetic variants with missing call rates exceeding 5% and samples with missing call rates exceeding 10% were removed. A minor allele frequency (MAF) threshold of >5% was applied to the overall dataset. In both sex-stratified and sex-combined modeling, the genotypes of female subjects were encoded as 0/1/2, which assumed entirely random X chromosome inactivation (XCI), and male genotypes were encoded as 0/2. [30] For sex-stratified analyses, the normalized expression was adjusted for covariates, including top 3 PCs, PEER factors [31], and sequencing platform. We also adjusted for sex in the sex-combined analysis. We fit linear regression models of expression against the covariates and kept the residuals as the adjusted expression. (Supplementary Figure 1) We obtained a list of 59 genes (Supplementary Table 1) that escape the XCI (XCI escapees) from Navarro-Cobos et al. [3] and investigated whether a different modeling strategy would improve the predictions of these escaping genes.

### Elastic net regression

We selected the elastic net regression to model gene expression levels on the X chromosome. While other approaches are available, elastic net regression allows for comparisons to both the autosomal models from PrediXcan [23] pipeline and published sex-stratified models of autosomal genes [32]. Elastic net regression is a regularized method that combines the LASSO (*L*_l_) and ridge (*L*_2_) penalties to select variables while fitting a linear regression model. Both penalty terms shrink regression coefficients by penalizing the regression model. The difference is that the LASSO penalty *L*_l_ often reduces some regression coefficients to 0 when the ridge penalty *L*_2_ does not.

The estimate from elastic net regression is 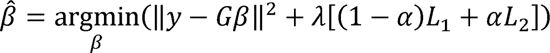, where y is the vector of the measured expression of one gene in all the individuals, and G is the genotype matrix of all the individuals. The genotypes of female subjects were encoded as 0/1/2, and male genotypes were encoded as 0/2. The shrinkage parameter A controls the amount of penalization applied to the regression. Theoretically, an elastic net model favoring the LASSO penalty (mixing parameter a= 0) will lead to a sparse model with a few SNPs with non-zero effects, while favoring the ridge penalty (mixing parameter α= 1) will lead to a polygenic model. We implemented k-fold nested cross-validation to tune each gene’s shrinkage parameter A at any given mixing parameter a and evaluate the model performance for how the model predicts on the hold-out fold. The best shrinkage parameter A was selected by the lowest mean squared error (MSE). The model assessment metrics include the coefficient of determination R^2^ and p-value, where the null hypothesis is that there is no correlation between predicted and measured expression. We only kept models with R> 0.1 and p-value < 0.05, the significant models, for the downstream analyses.

PrediXcan implements the elastic net regression model with a mixing parameter α = 0.5, where the penalty is defined as a balanced (50%-50%) combination of *L*_l_ and *L*_2_. This selection of α = 0.5 has been shown to be optimal by previous studies of autosomes. [23,33] Since we are unclear about the underlying regulatory function on the X chromosome, we fit elastic net regression models with different mixing parameters α = 0.05, 0.25, 0.5, 0.75, and 0.95. The models were fit in both sex-stratified and sex-combined manners. (Figure 1).

**Figure 1.**
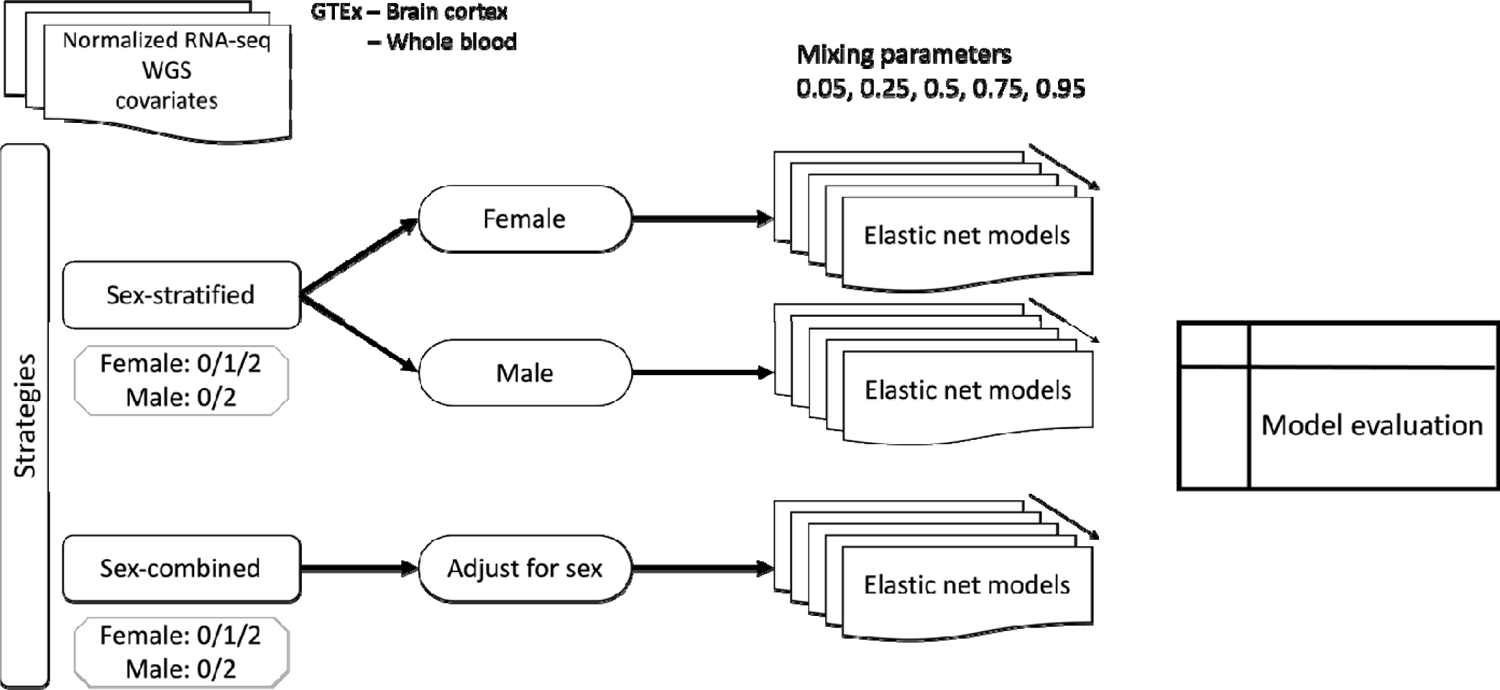
The working flow of model training using GTEx data. Elastic net regression models of the X chromosome gene expressions were trained in whole blood and brain cortex with different mixing parameters, in both sex-stratified and sex-combined manner. Genetic variants on the X chromosome were coded as 0/1/2 in female samples and 0/1 in male samples.

### Model comparisons with two-way ANOVA

We compared the performance of the fitted prediction models to determine (1) whether the widely adopted mixing parameter also works the best in X chromosome models; (2) whether sex-stratified models outperform the sex-combined models. Two-way ANOVA serves perfectly for our goal. Compared to one-way ANOVA, two-way ANOVA allows us to examine the combined effects of modeling strategies and biological factors.

We applied a data transformation to the model’s coefficient of determination to meet the normality assumption of two-way ANOVA since the distribution is right skewed with a long tail. The density plots and QQ plots of the original and the transformed are in Supplementary Figure 2. Though a light tail still exists, the transformed prediction performance metrics exhibit a more normal distribution than the original. The transformed shows homoscedasticity (homogeneity of variance) (see Supplementary Table 2). The design of GTEx and most other studies measuring human transcriptome provide independent observations of tissue-specific gene expression levels in each individual. Therefore, there is no violation of the assumptions of the two-way ANOVA test.

After transformation, we compare the model performance in different settings (mixing parameters, tissue types, female-vs. male-specific models, and sex-stratified vs. sex-combined analysis). We drew conclusions from the F statistics and p-values from the two-way ANOVA.

### Model Validation

We predicted the gene expression levels in the whole blood and brain cortex by applying the sex-stratified GTEx gene prediction models to DGN and MayoRNAseq data, respectively [23]. Genotypes of the DGN cohort were imputed to the HRC reference panel using the Michigan Imputation Server. We estimated the genetically regulated expression for each gene and tested its correlation with the measured expression with R^2^ statistics.

### AD risk analysis

We applied the validated significant models to sex-stratified GWAS meta-analysis summary statistics from the ADGC (Alzheimer’s Disease Genetics Consortium) with S-PrediXcan [34] pipeline, to identify potentially causal X-linked genes for AD. S-PrediXcan is a TWAS pipeline that allows summary statistics instead of individual-level data to impute the transcriptome and test for disease-associated genes. Due to the need for meta-analysis of the ADGC cohorts, we chose to perform this analysis using summary statistics rather than individual-level data. We matched the SNPs by rsIDs, since the models were trained in GRCh37 WGS data and the ADGC sex-stratified summary statistics are in GRCh38. A detailed description of the ascertainment of age at AD onset along with the descriptive statistics of each cohort can be found in Kunkle et al. [13]. All study participants were recruited under protocols approved by the appropriate institutional review boards. Written informed consent was obtained from study participants or, for those with substantial cognitive impairment, from a caregiver, legal guardian, or other proxy. The sex-stratified ADGC summary statistics detailed the effect of SNPs on clinically diagnosed AD in independent NHW subjects of the ADGC cohort. The summary statistics were from three different models stratified by sex: model 1 – additive effect of genetic variants adjusting for the first 3 PCs, model 2 – additive effect of genetic variants adjusting for the first 3 PCs and age, model 3 – additive effect of genetic variants adjusting for the first 3 PCs, age and APOE e4 allele count. These GWAS tests assumed no XCI. A total of 12,906 case participants (N=7,235 female cases) and 14,111 cognitively normal controls (N=8,411 female controls) were analyzed across 26 ADGC studies. Mean age-at-onset across studies ranged from 61.2 (standard deviation = 10.5) to 85.8 (SD = 5.7) and mean age-at-exam ranged from 64.0 (SD = 2.9) to 83.8 (SD = 7.5). Summary demographics of all studies can be found in Supplementary Table 3. The total number of SNPs analyzed for the male models ranged from 347,663 to 348,800 and for female models ranged from 487,238 to 489,735.

## Results

Given that there are numerous possible approaches to the analysis of X chromosome variants, we evaluated multiple options to determine the best modeling strategy for estimating the genetically regulated component of expression for genes on the X chromosome. We assessed whether the model parameters may differ by sex, and also attempted sex-combined and sex-stratified analyses. Given the unique biological features of the X chromosome, we also examined possible differences in the modeling of genes based on their X chromosome inactivation status. We further applied the significant models to AD summary statistics for a TWAS to identify AD risk genes on the X chromosome.

### Evaluation of mixing parameters within sex-stratified models

We trained sex-stratified models with GTEx data using five different mixing parameters: α = 0.05, 0.25, 0.5, 0.75, and 0.95. We used a nested cross-validation approach (similar to PrediXcan: https://github.com/hakyimlab/PredictDB-Tutorial) that tunes the shrinkage parameter A in an inner loop while model performance was evaluated in an outer loop. The models were fit in four tissue-sex sets: female-cortex, female-blood, male-cortex, and male-blood. We dropped intercept-only models since we are only interested in models that included genetic variants, and all the models will have at least one cis-eQTL with non-zero effect size on gene expression trait. As described in Methods, models with R< 0.1 and p> 0.05 were removed for downstream analyses.

Across different a, sex, and tissue types, we trained 499 significant gene models for 95 genes in whole blood and 135 genes in brain cortex. The average number of SNPs in each model is 21.3, and the average R^2^ = 0.12 (range from 0.03 to 0.54) (Table 2).

**Table 2.**
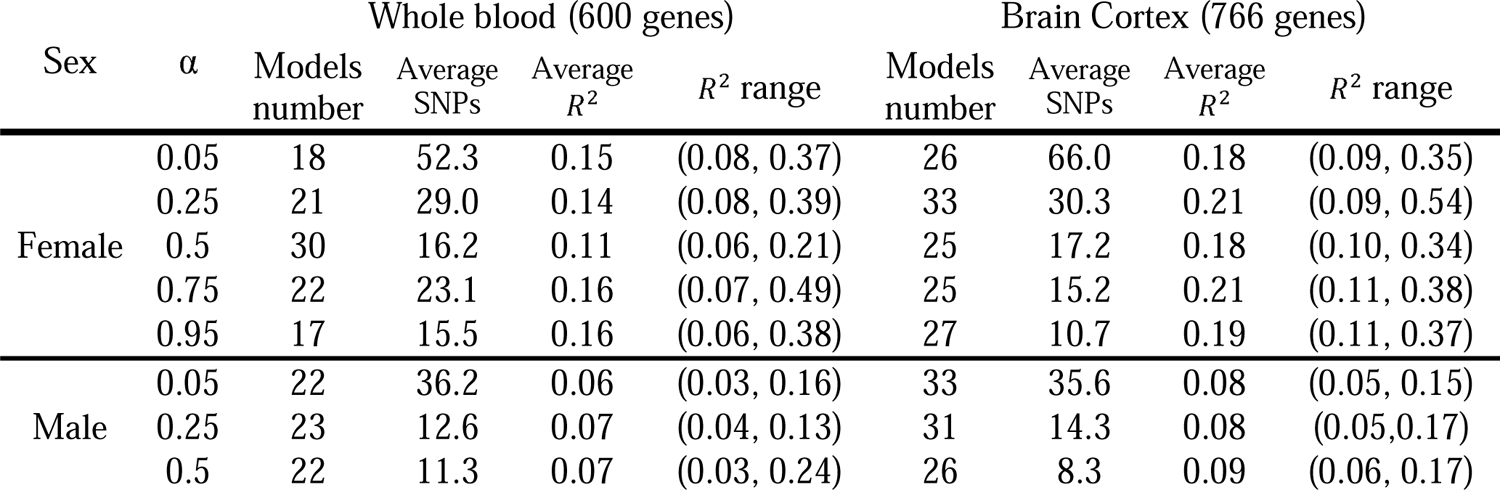
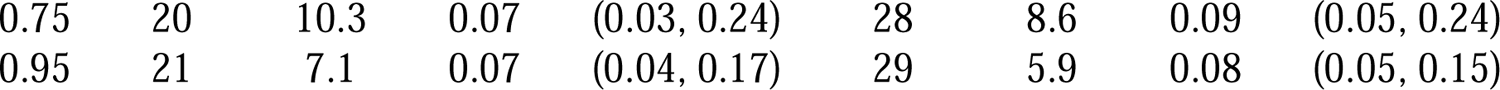
Summary of sex-specified models in blood and cortex with different mixing parameters. Only models with R> 0.1 and p < 0.05 were kept.

| Sex | $\alpha$ | Whole blood (600 genes) | | | | Brain Cortex (766 genes) | | | |
| --- | --- | --- | --- | --- | --- | --- | --- | --- | --- |
| | | Models number | Average SNPs | Average $R^2$ | $R^2$ range | Models number | Average SNPs | Average $R^2$ | $R^2$ range |
| Female | 0.05 | 18 | 52.3 | 0.15 | (0.08, 0.37) | 26 | 66.0 | 0.18 | (0.09, 0.35) |
|  | 0.25 | 21 | 29.0 | 0.14 | (0.08, 0.39) | 33 | 30.3 | 0.21 | (0.09, 0.54) |
|  | 0.5 | 30 | 16.2 | 0.11 | (0.06, 0.21) | 25 | 17.2 | 0.18 | (0.10, 0.34) |
|  | 0.75 | 22 | 23.1 | 0.16 | (0.07, 0.49) | 25 | 15.2 | 0.21 | (0.11, 0.38) |
|  | 0.95 | 17 | 15.5 | 0.16 | (0.06, 0.38) | 27 | 10.7 | 0.19 | (0.11, 0.37) |
| Male | 0.05 | 22 | 36.2 | 0.06 | (0.03, 0.16) | 33 | 35.6 | 0.08 | (0.05, 0.15) |
|  | 0.25 | 23 | 12.6 | 0.07 | (0.04, 0.13) | 31 | 14.3 | 0.08 | (0.05, 0.17) |
|  | 0.5 | 22 | 11.3 | 0.07 | (0.03, 0.24) | 26 | 8.3 | 0.09 | (0.06, 0.17) |

|  |  |  |  |  |  |  |  |  |
| --- | --- | --- | --- | --- | --- | --- | --- | --- |
| 0.75 | 20 | 10.3 | 0.07 | (0.03, 0.24) | 28 | 8.6 | 0.09 | (0.05, 0.24) |
| 0.95 | 21 | 7.1 | 0.07 | (0.04, 0.17) | 29 | 5.9 | 0.08 | (0.05, 0.15) |

In both whole blood and brain cortex, the transformed R^2^ of female-only models present a wider spread and higher central tendency than male-only models (Supplementary Figure 3). We checked the distributions of the original R^2^in the sex-stratified models. For each tissue-sex set, we compared the original R^2^of the significant gene models with the R^2^of the same genes in the other tissue-sex set. (See Figure 2 for the comparison of α= 0.5 and Supplementary Figure 4-7 for the other mixing parameters)

**Figure 2.**
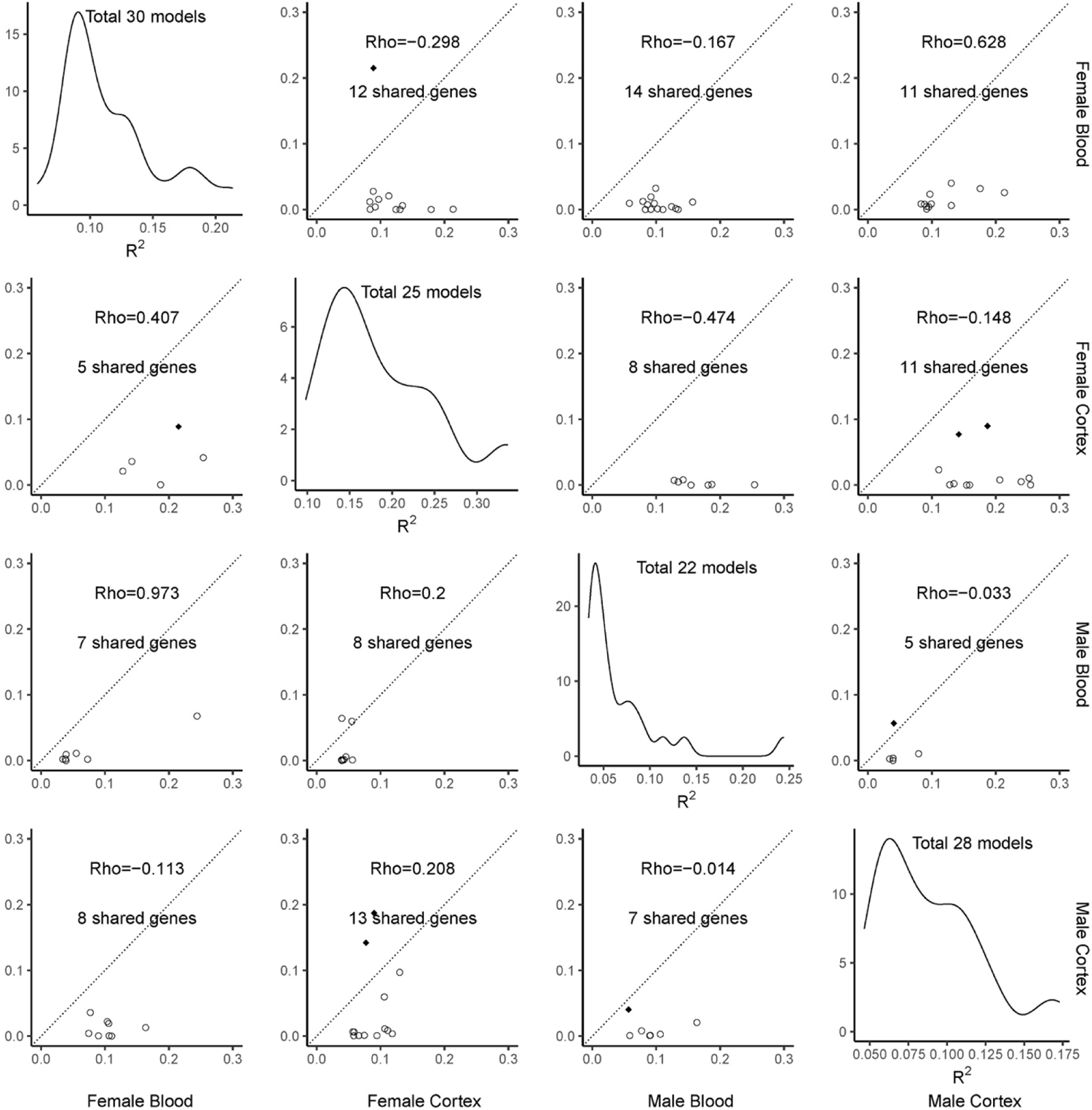
Comparison matrix of the balanced sex-stratified elastic net models trained in GTEx whole blood and brain cortex data (α = 0.5). The four diagonal panels are the distributions of model R^2^. The number of genes being modeled in each tissue-sex set is listed in the distribution plot. The off-diagonal panels compared the R^2^ of the models in pairs of tissue-sex sets. The four rows are for the female-blood, female-cortex, male-blood, and male-cortex. For each row, the genes of the significant models in the tissue-sex set were matched in the other sets for the matching R^2^. The diamonds represent genes with a significant model (R < 0.1 and p > 0.05) in another tissue-sex set, while the circles represent genes with a non-significant model in another tissue-sex set. The number of gene models and the Pearson correlation coefficients were listed in the comparison plots

We performed a two-way ANOVA under the null hypothesis that there are no differences in the transformed R^2^ among sex-stratified elastic net models with different mixing parameters α = 0.05, 0.25, 0.5, 0.75, and 0.95. In both cortex and whole blood models, we found no significant differences in the average R^2^ among different a’s in sex-stratified models (p = 0.21 for blood, p = 0.55 for cortex), but did observe significant differences in the average R^2^ by the sample sex (p = 1.58 × 10^-26^ for blood, p = 1.83 × 10^-55^for cortex). This result is driven by lower average R^2^in male-only models conditioned on tissue and a, as shown in Table 2. Sex by a interaction was also non-significant (p = 0.69 for blood, p= 0.55 for cortex). This finding indicates that different mixing parameters generally do not impact the performance of gene expression prediction models. In contrast, the models trained in female- and male-only samples have different coefficients of determination. While our elastic net models capture more genes with α = 0.25, to keep consistent with the literature and existing PredictDB autosomal models, we used a mixing parameter of α = 0.5 (balanced penalty) for all subsequent analyses.

### Differences in the tissue-specific sex-stratified models

For 103 models with a mixing parameter of α= 0.5, the average number of SNPs in each model is 13.4, and the average R^2^ = 0.116 (range from 0.03 to 0.34). There are 30, 22, 25, and 26 significant models for female-only blood, male-only blood, female-only cortex, and male-only cortex samples, respectively (see Figure 2). Among all the genes modeled, no blood-specific expression was modeled both in males and females, while two cortex-specific expressions (ENSG00000005893 *LAMP2*, ENSG00000101955 *SRPX*) were modeled in both sexes, and one gene was modeled in both tissues in male- or female-only data (ENSG00000197582 *GPX1P1*, ENSG00000013563 *DNASE1L1*).

We further compared the differences between models trained with data from different tissue types. We found a statistically significant difference in the model’s average R^2^by sex (p= 1.93 × 10^-l3^). As expected, we observed a statistically significant difference between whole blood and brain cortex (p = 5.33 × 10^-8^). As shown in Table 2, the average R^2^of blood-specific models is lower than cortex-specific models, and the average R^2^ of male-only models is lower than female-only models. Notably, there were no significant differences noted between sex-stratified models of autosomal genes from the same dataset (reported in Table 1 of Mahoney et al. [32]), regardless of sample size differences between males and females. The interaction between sex and tissue, however, was not significant (p= 0.59). We highlight that sex differences are just as crucial as tissue differences for X chromosome gene expression.

### Sex-combined models with a balanced penalty

The tissue-specific sex-combined models were trained in blood and cortex, respectively, with a balanced mixing parameter a = 0.5. After removing intercept-only models and models with R< 0.1 and p > 0.05, we have 45 significant sex-combined models in whole blood and cortex. The average number of SNPs in the sex-combined models across different tissues is 10, while the average R^2^ = 0.052 (range from 0.023 to 0.11). (Table 3) The average R^2^ of the sex-combined models is smaller than that of sex-stratified models (single-sided t-test p < 2.2 × 10^-l6^).

**Table 3.** Summary of sex-combine, balanced penalty models in blood and cortex. Models with R< 0.1 and p > 0.05 were removed. The models are summarized conditioned on the XCI escaping status. None of the genes escaping XCI were significant in blood.

| Tissue | Escape XCI | Models number | Average SNPs | Average $R^2$ | $R^2$ range |
| --- | --- | --- | --- | --- | --- |
| Blood | × | 23 | 8.96 | 0.045 | (0.023, 0.087) |
| Cortex | × | 20 | 11 | 0.059 | (0.03, 0.11) |
|  | √ | 2 | 11.5 | 0.051 | (0.034, 0.067) |

We compared different modeling strategies (stratified-female, stratified-male, and sex-combined models) to determine whether the sex-combined analysis would improve the overall fit for X chromosome genes by increasing the sample size. Considering the potential different underlying regulatory effects, we tested the genes escaping X chromosome inactivation (XCI escapees) and the other genes separately. Besides, there are some XCI escapees in the sex-stratified balanced penalty models (3 for female-blood, 2 for male-blood, and 1 for male-cortex).

For the models of XCI escapees (n=8), we found no significant differences in the average R^2^ in the modeling strategies (p = 0.15) nor tissue types (p= 0.55). Contrarily, in both blood and cortex, genes not escaping from the XCI show significantly different R^2^ between any pair of modeling strategies (one-way ANOVA p= 3.4 × 10^-ll^ for blood and p = 1.01 × 10^-l5^ for cortex).

### Model Validation

The sex-stratified GTEx gene prediction models in blood and cortex were applied to DGN and MayoRNAseq validation datasets, respectively. 65 SNPs from GTEx whole blood models were not available in DGN whole-genome imputed genotype data, and 15 SNPs from GTEx brain cortex models were not available in MayoRNAseq WGS data (Figure 3 A and B). In DGN whole blood validation, 20 and 15 genes were predicted by stratified-female and stratified-male significant models, respectively. There is no gene predicted in both sexes. In MayoRNAseq TCX validation, we predicted 19 and 18 genes with stratified-female and stratified-male significant models, respectively, among which two genes were predicted in both females and males. (Figure 3 C and D). In addition, two sex-combined significant models were applied to genes that escape from XCI in MayoRNAseq data.

**Figure 3.**
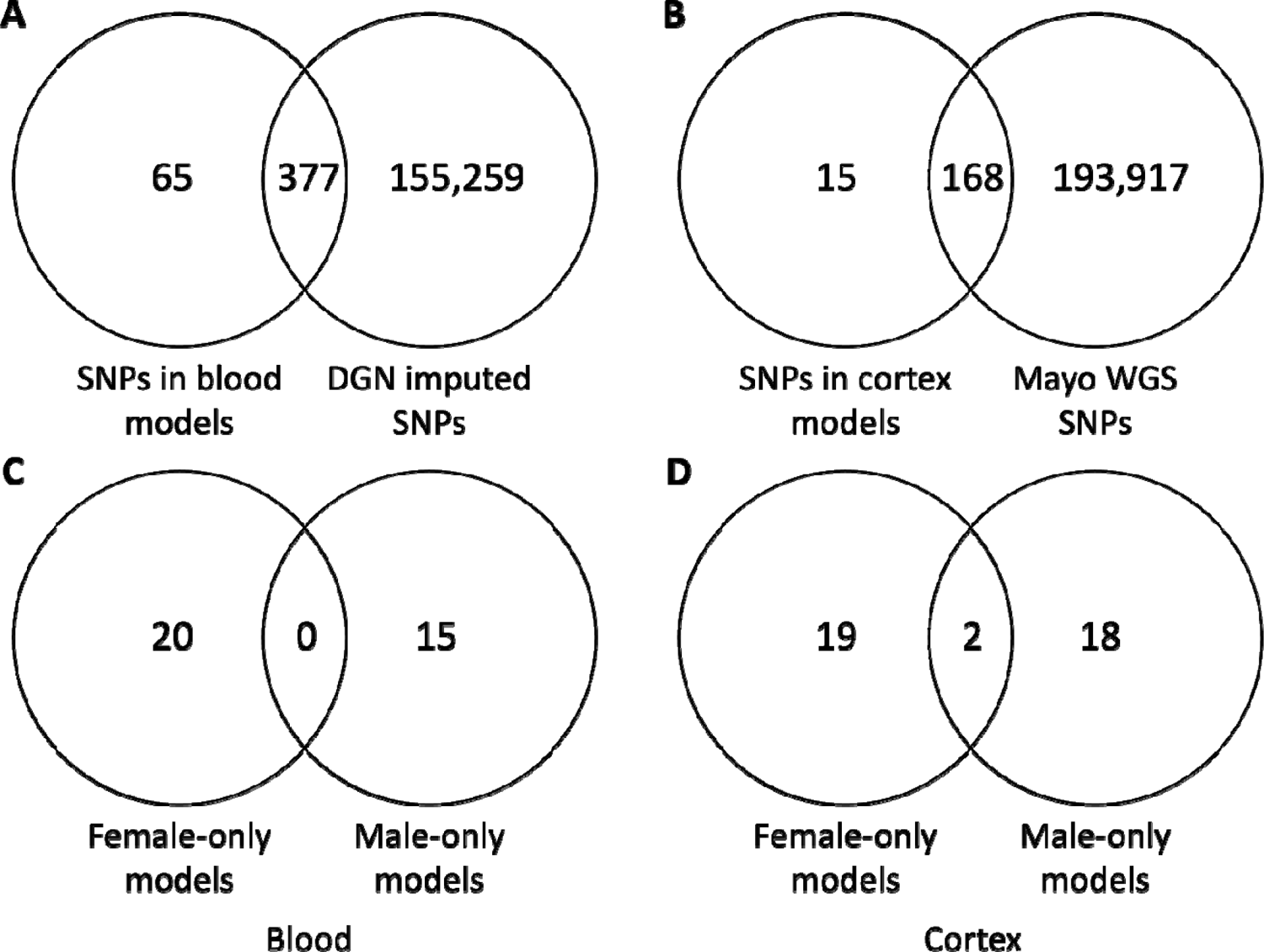
The number of SNPs in the validation datasets and the number of genes predicted. A and B: Venn diagrams representing the number of SNPs in the tissue-matching validation datasets and our sex-stratified prediction models. C and D: Venn diagrams illustrating the number of genes predicted by the sex of the models and validation samples

We calculated prediction R^2^ by regressing the estimated expressions against the measured expression via RNA-seq from the validation datasets (range from 9.94 × 10^-5^ to 0.091). Table 4 lists the prediction R^2^ of the sex-stratified and sex-combined models in each tissue. Comparisons of the expected and observed distribution of prediction R^2^ in blood samples (DGN) and cortex samples (MayoRNAseq) for all models (Supplementary Figure 8) show a deviation from the expected null distribution, demonstrating that models generally explain variability in X chromosome gene expression.

**Table 4.** Prediction R^2^ in validation datasets. Note that there is no significant blood-specific sex-combined models for XCI escapees.

|  | Blood – DGN |  |  | Cortex – MayoRNAseq |  |  |
| --- | --- | --- | --- | --- | --- | --- |
| | Models Number | Average $R^2$ | $R^2$ range | Models Number | Average $R^2$ | $R^2$ range |
| Female-only model | 20 | 0.014 | $(9.99 \times 10^{-6}, 0.074)$ | 21 | 0.0097 | $(9.94 \times 10^{-5}, 0.091)$ |
| Male-only model | 15 | 0.034 | $(0, 0.44)$ | 20 | 0.010 | $(1.05 \times 10^{-4}, 0.036)$ |
| Sex-combined model (Only XCI escapees) | - | - | - | 2 | 0.0010 | $(1.28 \times 10^{-4}, 0.0020)$ |

### Identify AD risk genes on X chromosome

We applied sex-stratified gene prediction models with a balanced penalty to the ADGC sex-stratified GWAS summary statistics on the X chromosome for TWAS analysis, using the S-PrediXcan [34] pipeline. In this sex-stratified analysis of X-linked AD risk genes, 25 and 26 genes were tested in cortex in females and males respectively, while 30 and 22 genes were tested in blood in females and males respectively. We found a marginally significant gene *ARMCX6* in cortex in female-only data (p = 0.0082, while the Bonferroni correction significance threshold is 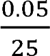 = 0.002). Other associations we found from this TWAS are presented in Table 5.

**Table 5.** S-PrediXcan results (p<0.05) on sex-stratified AD summary statistics.

| Sex | GWAS model | Tissue | Gene Name | Z score | p-value | Number of SNPs used |
| --- | --- | --- | --- | --- | --- | --- |
| Female | Additive + Age + APOE | Cortex | *ARMCX6* | 2.64 | 0.0082 | 23 |
| Male | Additive + Age + APOE | Cortex | *DIAPH2* | -2.31 | 0.0211 | 15 |
| Female | Additive | Blood | *LL0XNC01-116E7.2* | 2.3 | 0.0217 | 17 |
| Female | Additive + Age | Blood | *TENM1* | -2.26 | 0.0238 | 52 |
| Male | Additive | Blood | *SUV39H1* | -2.16 | 0.0308 | 2 |
| Female | Additive | Blood | *TENM1* | -2.15 | 0.0316 | 52 |
| Male | Additive + Age | Blood | *SUV39H1* | -2.05 | 0.0407 | 2 |
| Female | Additive + Age | Cortex | *ARMCX6* | 2.03 | 0.0426 | 23 |
| Female | Additive + Age | Cortex | *LAMP2* | 2.02 | 0.0435 | 20 |
| Female | Additive + Age | Blood | *LL0XNC01-116E7.2* | 2.01 | 0.0444 | 17 |
| Female | Additive | Cortex | *UBQLN2* | 2.01 | 0.0446 | 5 |
| Male | Additive + Age | Cortex | *TMSB4X* | -2.01 | 0.0447 | 2 |
| Female | Additive | Cortex | *ARMCX6* | 2 | 0.0454 | 23 |
| Female | Additive | Cortex | *LAMP2* | 2 | 0.0456 | 20 |
| Male | Additive + Age + APOE | Blood | *TBC1D25* | 1.99 | 0.0465 | 1 |
| Female | Additive | Cortex | *DNASE1L1* | -1.96 | 0.0494 | 8 |

The prediction model of *ARMCX6* in brain cortex, trained with female-only data, selected 23 SNPs from the 164 SNPs in the *cis*-regulatory region. The shrinkage parameter A of the model is 0.25. From 5-fold cross-validation, the correlation between predicted and observed expression levels is R^2^ = 0.333 (p= 1.3 × 10^-4^). This gene model was also trained in male-only cortex data, but it is not significant (R^2^ = 0.01, p = 0.39), while no model of *ARMCX6* from blood data exists. Based on our TWAS results, increases in the genetically predicted expression of *ARMCX6* in cortex will increase the AD risk of a female subject (p = 0.0082).

We visualized the putative AD-risk gene, *ARMCX6* in Figure 4. All the SNPs were plotted in the GRCh37 positions to make them comparable across panels. The positions and weights of the 23 SNPs included in the *ARMCX6* female-only penalized prediction model are presented in Figure 4A. Only 2 SNPs are located upstream of the *ARMCX6* gene. The remaining 21 SNPs form multiple clusters localized to 1,000 bp windows downstream of the *ARMCX6* gene. Figure 4B and Figure 4C show the p-values of SNPs within the same region (as Figure 4A) from female-only eQTL mapping and female-only GWAS on AD (adjusted for age and APOE), respectively. The SNPs in the prediction model are in red. None of the selected SNPs are of the most extreme eQTL or GWAS p-values. The female-only eQTL p-values were calculated with GTEx NHW female subjects with brain cortex samples (N=46). The distributions of *ARMCX6* gene expression in both sexes in brain cortex and blood from GTEx are in Supplementary Figure 9.

**Figure 4.**
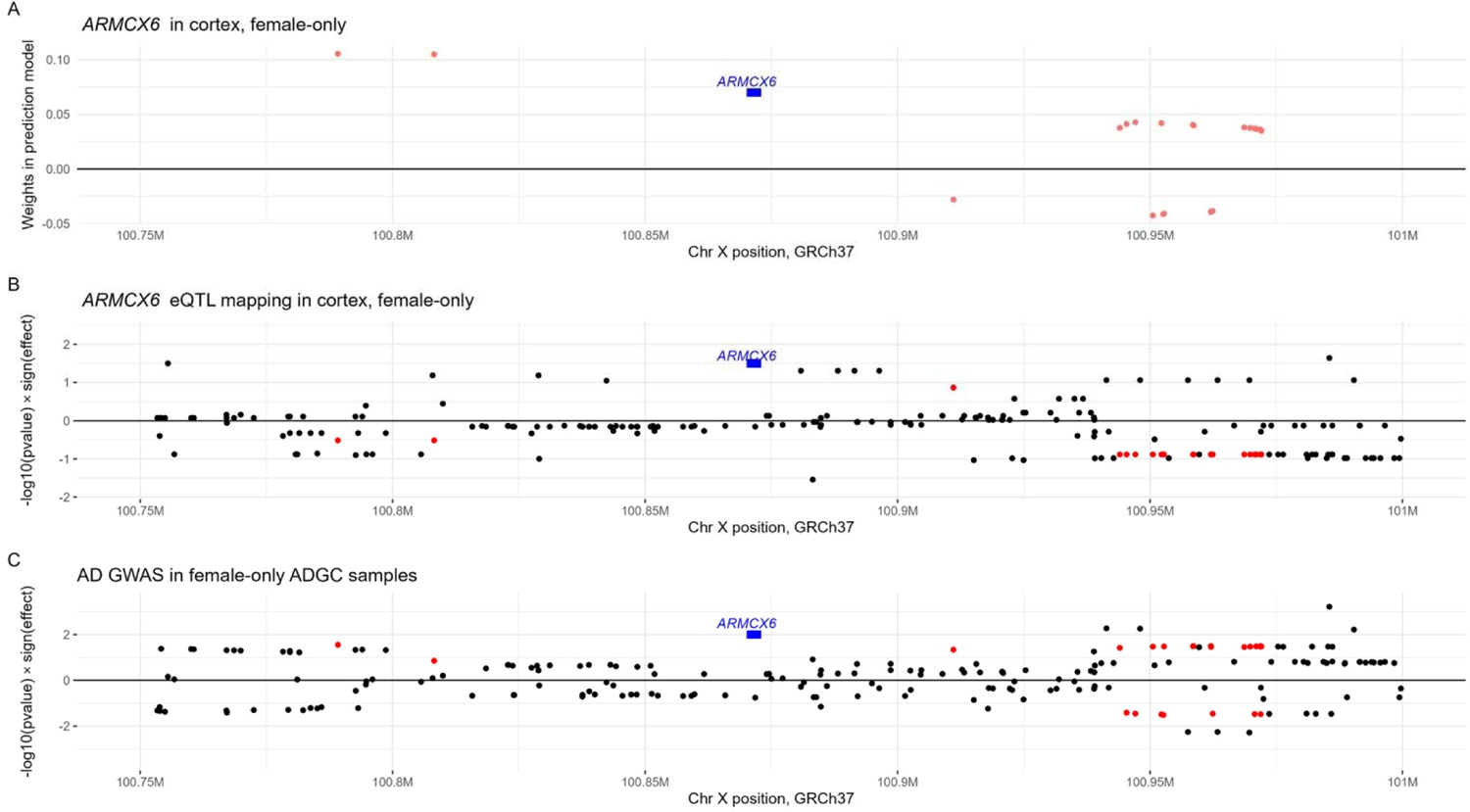
Genetic architecture of TWAS significant hit, ARMCX6. All three panels show the same region. A: The positions and effect sizes of the 23 selected SNPs in the cortex-specific expression prediction model trained with female-only samples. B: Hudson plot of eQTLs of ARMCX6 in female-only GTEx cortex samples. C: Hudson plot of ADGC meta-analysis GWAS in female-only samples. The red dots in B and C represent the SNPs in the expression prediction model

One-way ANOVA suggests the expression levels vary among the four tissue-sex sets (p= 0.014), while two-way ANOVA indicates the differences come from tissue (p = 0.0062) but not sex (p = 0.18).

## Discussion

In this work, we examined multiple modeling strategies for generating X chromosome gene expression prediction models. Based on our cross-validation results from analyses of the GTEx data for whole blood and brain cortex, we concluded that sex-stratified models (with a= 0.5) provided the optimal R^2^. We validated these models in two external tissue-matched datasets and found an average prediction performance of 0.012, which deviates from the expected distribution under a null hypothesis. With these models, we implemented a TWAS and tested the association between AD risk and 20 genes for females and 15 genes for males on the X chromosome in whole blood, 21 genes for females and 20 genes for males on the X chromosome in brain cortex. *ARMCX6* is our most compelling finding, as we did not find any gene significant after strict Bonferroni correction.

DNA methylation silences one copy of X chromosome genes in females via X inactivation, and there are noted non-random patterns associated with this phenomenon, particularly skewed [6,7] and escaped [2,3] X inactivation. We were surprised to observe that modeling genes escaping XCI with different strategies produced no substantial differences compared to non-escaped genes. In our sex-stratified models, we noted that our modeling approach assumes equal impact of each allele, which for the X chromosome would also assume totally random X chromosome inactivation. Accounting for skewed X-inactivation may be possible with availability of DNA methylation data or allele-specific expression analyses.

We note that the heterogeneity in gene expression prediction between males and females was substantial, and that these differences were not overcome by the power gain from increasing the sample size when combining males and females. Our X chromosome gene models fit within males and females are remarkably different from prior reports of similar sex-stratified analyses of the autosomes, also of the GTEx dataset [32]. This finding provides support that differences between male and female models are not driven by sample size differences and may also point to biological differences in the evolution of the X chromosome. For example, prior reports have shown depletion of regulatory variants on the X chromosome [20], etc.

Our strategy of selecting *statistically significant* expression prediction models only is conservative, and the general distribution of prediction R^2^ suggests that models not meeting our statistical significance criteria may still be useful in TWAS predictions. We also chose an a = 0.5 for all subsequent analyses as the R^2^ distribution was not significantly different from our ANOVA analyses; however, we note that in cortex specifically, slightly more genes are modeled using an a = 0.25. Therefore, for some TWAS applications, a lower alpha threshold may be preferred to capture additional genes.

Even though we only reported the elastic net penalized models in brain cortex and blood, the modeling strategy can be applied to all the other tissues to train the tissue-specific gene expression prediction models. It will then be possible to detect putative sex-specific disease risk genes on the X chromosome with individual level genetic data or GWAS summary statistics.

*ARMCX6*, as our putative AD risk gene, also has nominally significant p-values (p<0.05) when the GWAS model did not adjust for APOE (p = 0.0426) or age (p = 0.0454). Due to the correlation among gene expression models by tissue and sex, a Bonferroni correction for all statistical tests is overly conservative; we, therefore, corrected our hypothesis tests for 25 tests (in female-only cortex TWAS) to establish an α = 0.002.

*ARMCX6* was reported to be associated with mitochondrial dynamics in neurons [35]. *ARMCX6* and other genes in the *Armcx* cluster are highly expressed in the human nervous system. These genes encode a family of mitochondrial proteins regulating mitochondrial trafficking, which may increase the risk of neurodegenerative diseases. A human brain proteomics study [36] suggests that *ARMCX6* has sex-differentiated expression at the mRNA level.

There are a few SNPs near *ARMCX6* that were reported to be GWAS hits from the GWAS Catalog[37]. None of these variants were selected in our modeling process; some variants were not sequenced in GTEx, while others were removed by the MAF threshold. One GWAS hit, rs5951278, 25,389 base pairs upstream of *ARMCX6* is associated with educational acquisition [38]. This association on the X chromosome can either represent education as a potential confounder of the protection from AD or suggest a potentially shared pathway between neural development/degeneration and educational attainment. In addition, rs148260947, 271,216 base pairs upstream of *ARMCX6*, is reported to be associated with cognitive function measured by mini-mental state examination (MMSE) at baseline [18].

The TWAS Atlas [39] reports two X chromosome genes associated to AD, *ZC3H12B* [40] and *ARHGAP4* [41]. We could not generate reliable sex-stratified TWAS models for either of these genes in our analyses. Notably, these prior studies applied TWAS pipelines to the X chromosome without addressing genetic differences due to sex. We have demonstrated from our models that gene expression on the X chromosome should be studied in a sex-stratified manner.

There are several limitations to our study. First, we have a limited sample size; while the sex- combined eQTL sample sizes are generally substantially smaller than other types of genetic studies due to the expected larger effect sizes, our sex-stratified analyses are effectively halving the total sample. This extremely small sample size on the X chromosome limits the power for both eQTL mapping and prediction modeling. We also restricted analyses to whole blood and cortex tissues in this study. This was due to the availability of validation datasets and their relevance for AD. Many AD GWAS datasets and eQTL datasets of brain samples have excluded genetic variation on the X chromosome, a practice which persists despite attempts by funding agencies to promote sex chromosome studies. Our small X chromosome eQTL sample size limited the power to detect AD risk genes with TWAS pipelines. Another potential approach to allow for sex-specific effects while conserving statistical power is meta-analysis [21,22].

Therefore, we explored the possibility of performing a meta-analysis across sex-stratified models in each tissue using the LASSOSum approach [42]; however, these models were difficult to fit due to model convergence issues and were not included in our evaluations.

Despite these limitations, this work provides a set of prediction models to enable X chromosome TWAS analyses of blood and cortex. Based on our results, we recommend using sex-stratified models for all X chromosome genes. We also detect a putative risk gene, *ARMCX6*, for AD on the X chromosome, which highlights the potential role of X chromosome variation in Alzheimer’s disease.

## Acknowledgments

This work made use of the High Performance Computing Resource in the Core Facility for Advanced Research Computing at Case Western Reserve University. Special thanks to Makaela Mews for the sex-stratified meta-analysis with LASSOSUM, as well as Yousef Mustafa’s insight into visualizing TWAS results. The Genotype-Tissue Expression (GTEx) Project was supported by the Common Fund of the Office of the Director of the National Institutes of Health, and by NCI, NHGRI, NHLBI, NIDA, NIMH, and NINDS. The data used for the analyses described in this manuscript was obtained from dbGaP accession number phs000424.v7.p2 on 04/03/2018. The results published here are in part based on the MayoRNAseq data obtained from the AD Knowledge Portal (https://adknowledgeportal.org). The data available in the AD Knowledge Portal would not be possible without the participation of research volunteers and the contribution of data by collaborating researchers.

## Availability statement

The GTEx data is openly available via from dbGaP. The MayoRNAseq data is openly available at the AD Knowledge Portal (https://adknowledgeportal.org). The sex-stratified predictive models of X chromosome genes in whole blood and brain cortex are available at https://github.com/bushlab-genomics/ChrX_predixcan.

