## Supplementary Figure for "An X Chromosome Transcriptome Wide Association Study Implicates ARMCX6 in Alzheimer’s Disease"

### Supplementary Figures

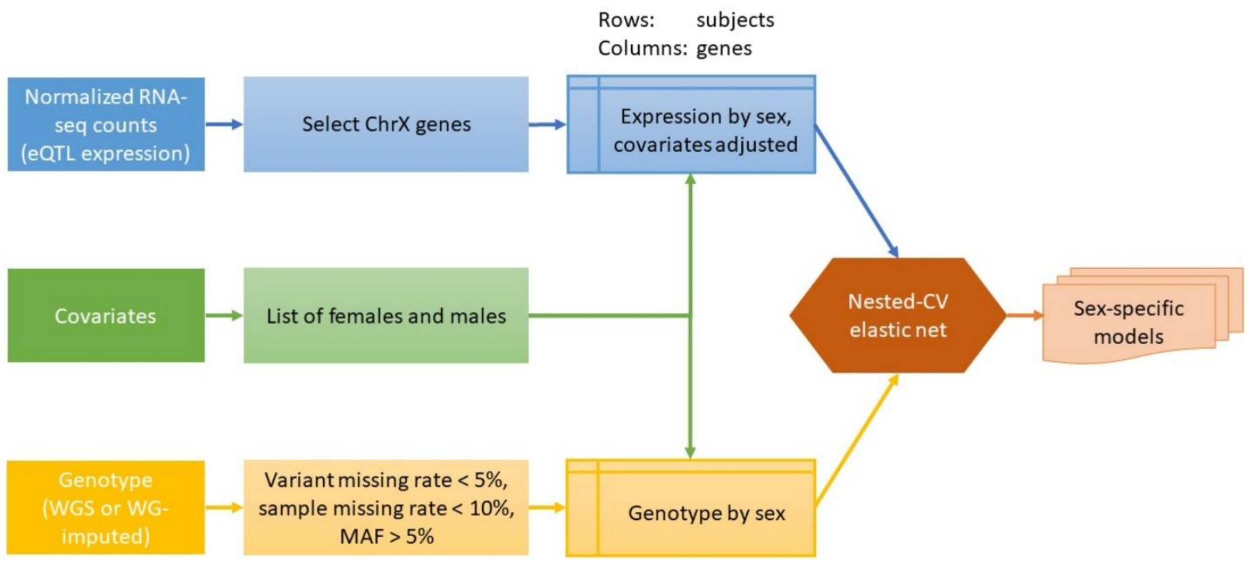


*Supplementary Figure 1. Data manipulation and quality control.*

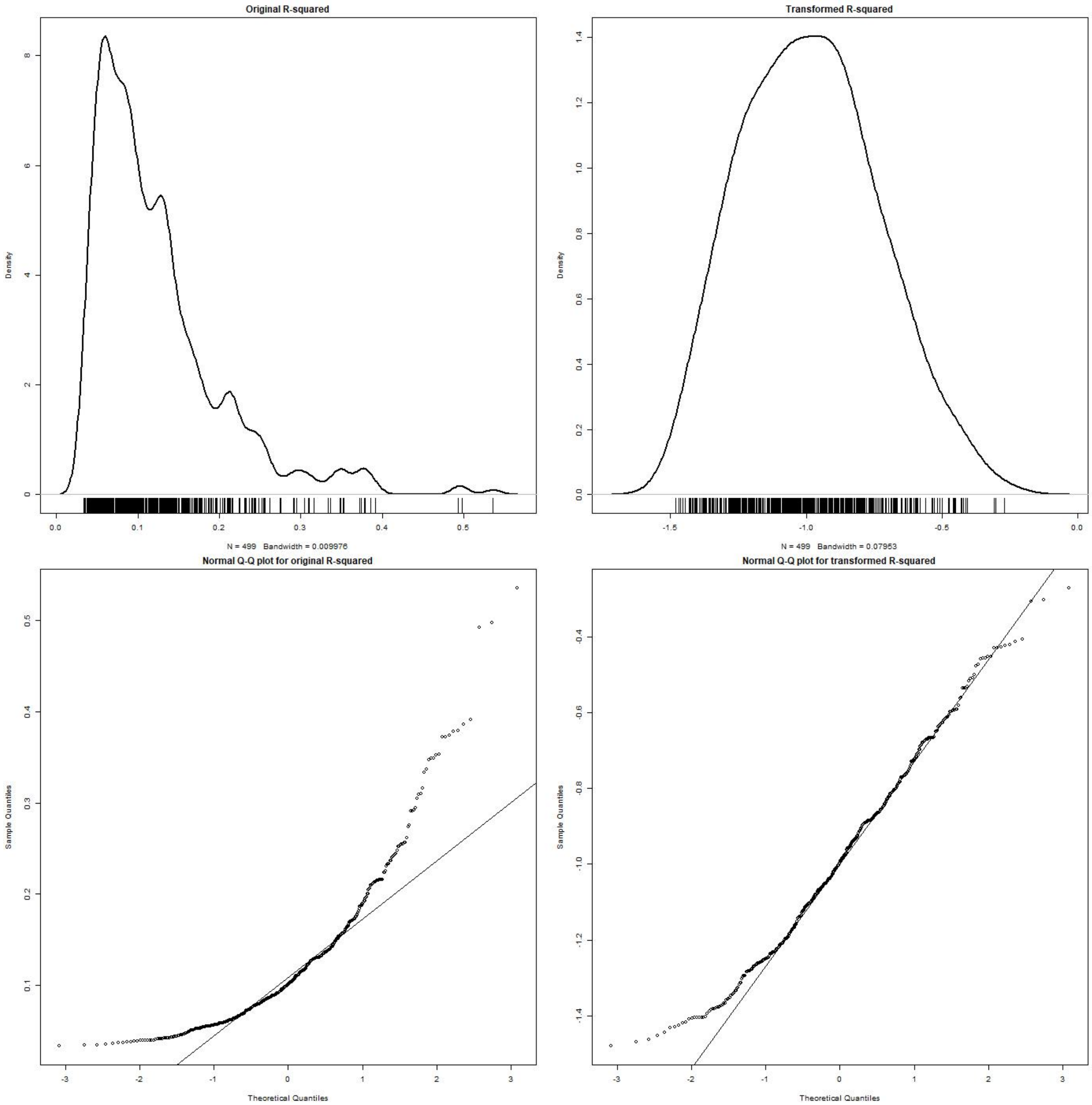


*Supplementary Figure 2. Density distribution and QQ plot of all models' $R^2$ before and after transformation.*

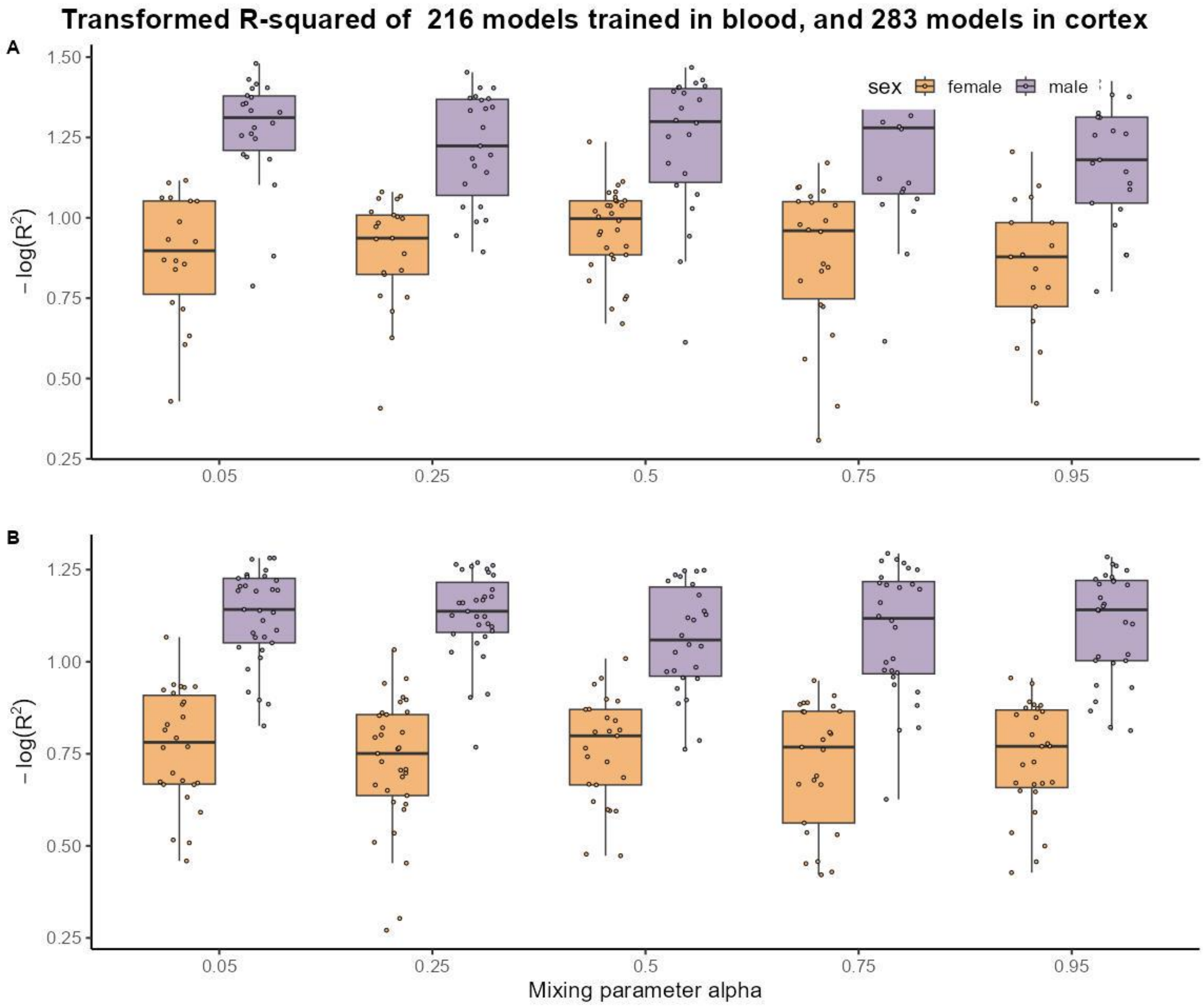


*Supplementary Figure 3. Boxplots of the transformed $R^2$ sex-stratified models in whole blood (A) and in brain cortex (B)*

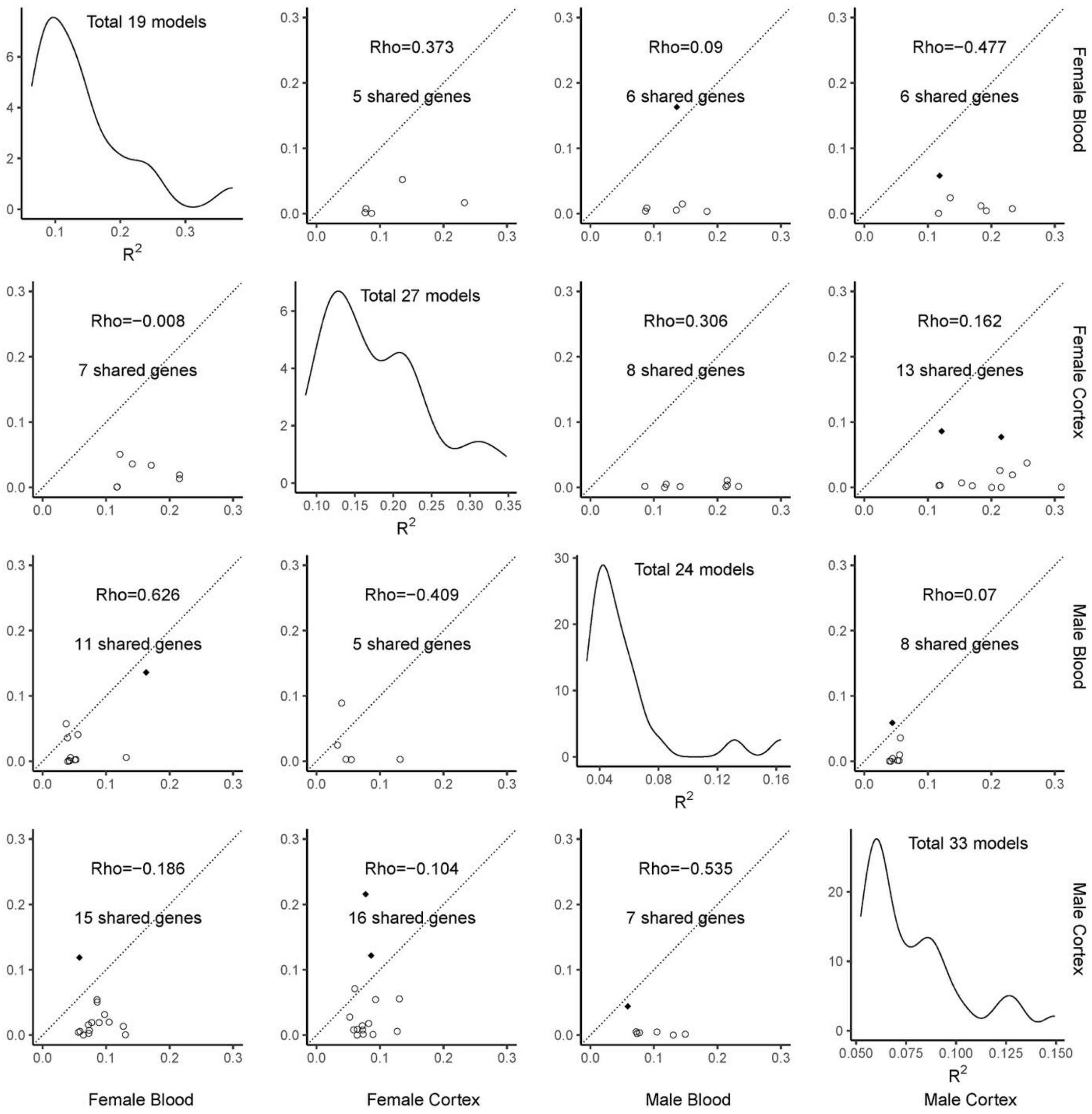


*Supplementary Figure 4. Comparison matrix of the balanced sex-stratified elastic net models trained in GTEx cortex and blood data, α=0.05. The four diagonal panels are the distributions of model $R^2$. The number of genes being modeled in each tissue-sex set is listed in the distribution plot. The off-diagonal panels compared the $R^2$ of the models in pairs of tissue-sex sets. The four rows are for the female-blood, female-cortex, male-blood, and male-cortex. For each row, the genes of the significant models in the tissue-sex set were matched in the other sets for the matching $R^2$. The diamonds represent genes with a significant model ($R < 0.1$ and $p > 0.05$) in another tissue-sex set, while the circles represent genes with a non-significant model in another tissue-sex set. The number of gene models and the Pearson correlation coefficients were listed in the comparison plots.*

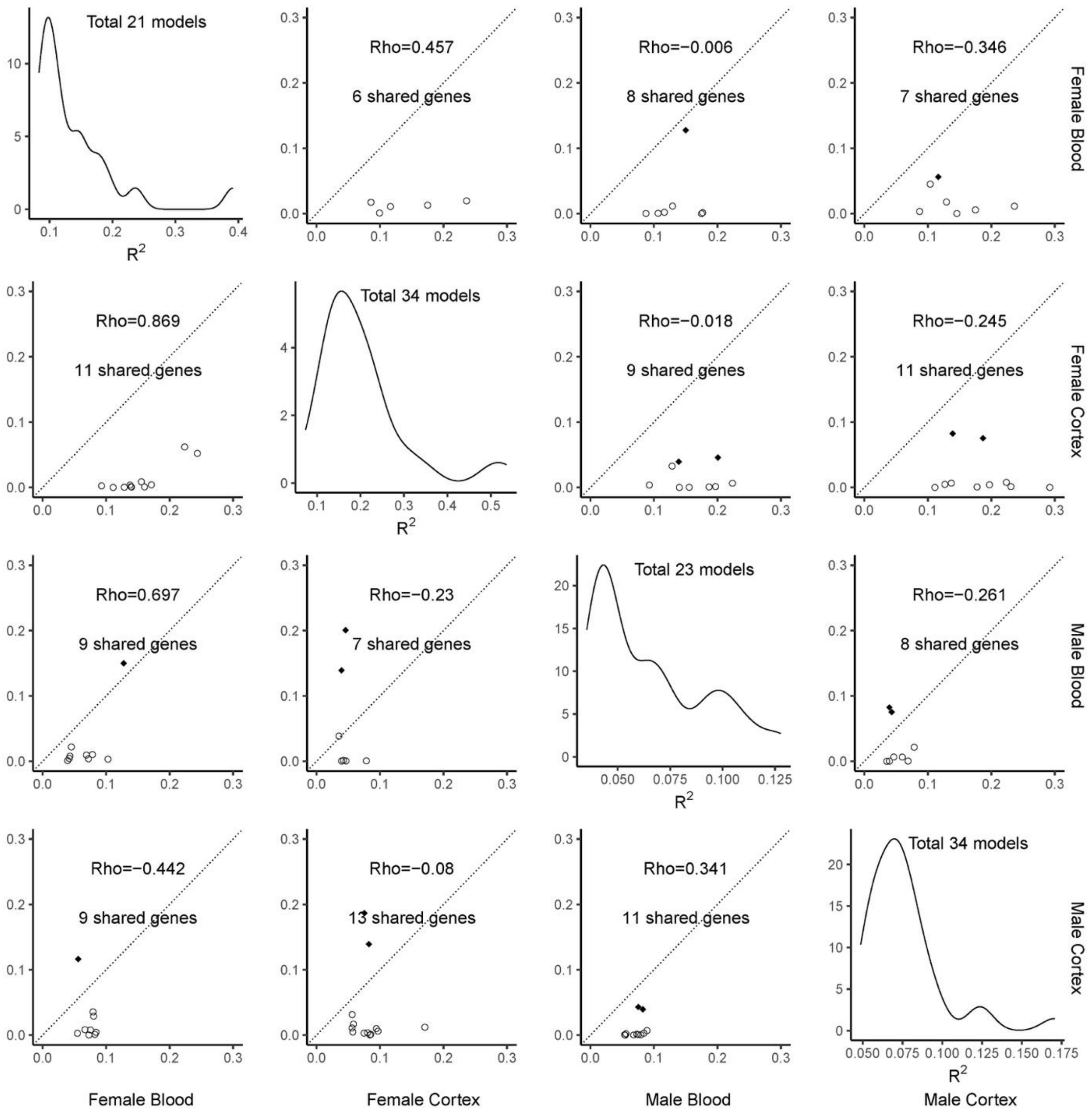


*Supplementary Figure 5. Comparison matrix of the balanced sex-stratified elastic net models trained in GTEx cortex and blood data, α=0.25. The number of genes being modeled in each tissue-sex set is listed in the distribution plot. The off-diagonal panels compared the $R^2$ of the models in pairs of tissue-sex sets. The four rows are for the female-blood, female-cortex, male-blood, and male-cortex. For each row, the genes of the significant models in the tissue-sex set were matched in the other sets for the matching $R^2$. The diamonds represent genes with a significant model ($R < 0.1$ and $p > 0.05$) in another tissue-sex set, while the circles represent genes with a non-significant model in another tissue-sex set. The number of gene models and the Pearson correlation coefficients were listed in the comparison plots.*

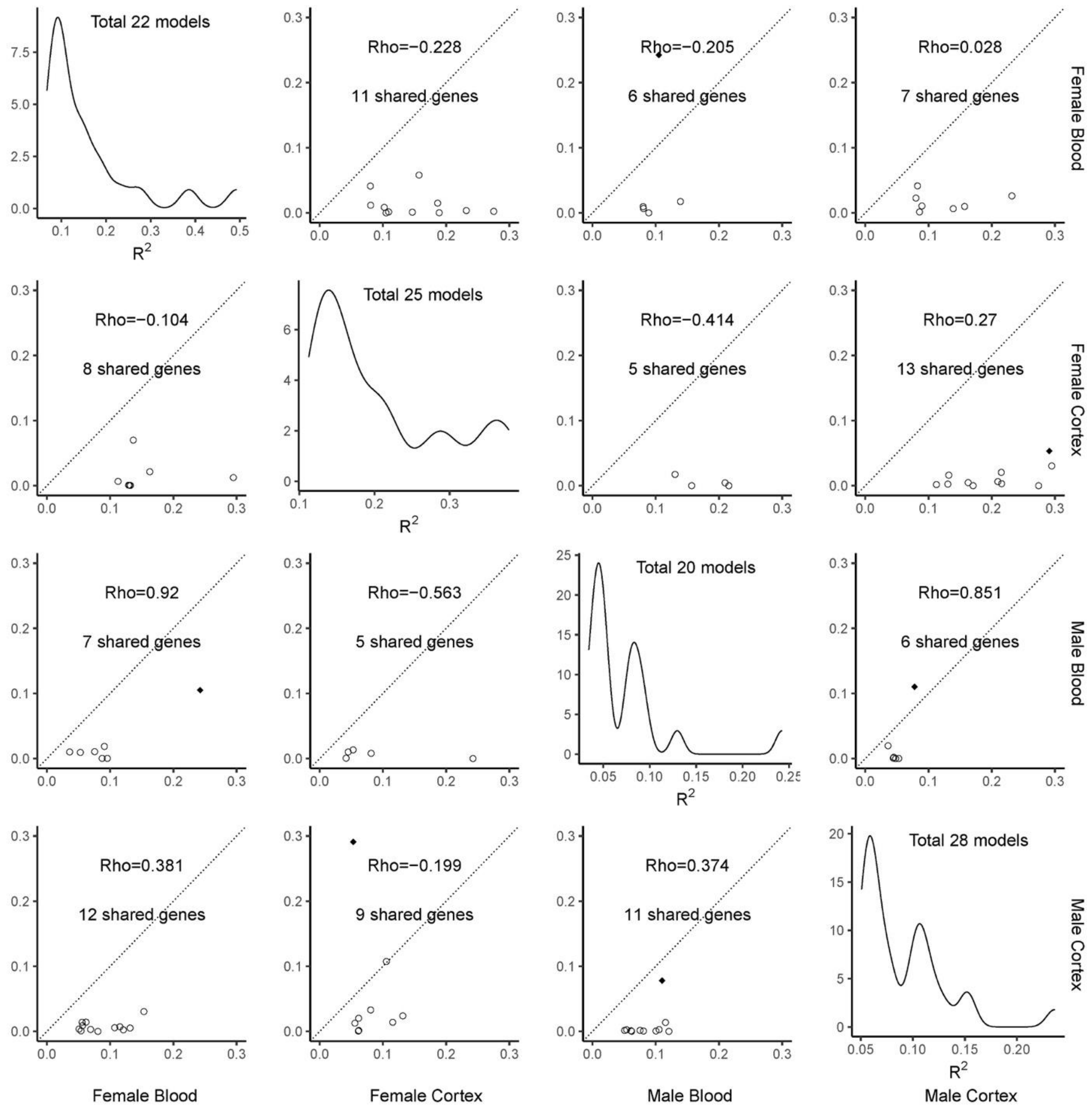


*Supplementary Figure 6. Comparison matrix of the balanced sex-stratified elastic net models trained in GTEx cortex and blood data, α=0.75. The number of genes being modeled in each tissue-sex set is listed in the distribution plot. The off-diagonal panels compared the $R^2$ of the models in pairs of tissue-sex sets. The four rows are for the female-blood, female-cortex, male-blood, and male-cortex. For each row, the genes of the significant models in the tissue-sex set were matched in the other sets for the matching $R^2$. The diamonds represent genes with a significant model ($R < 0.1$ and $p > 0.05$) in another tissue-sex set, while the circles represent genes with a non-significant model in another tissue-sex set. The number of gene models and the Pearson correlation coefficients were listed in the comparison plots.*

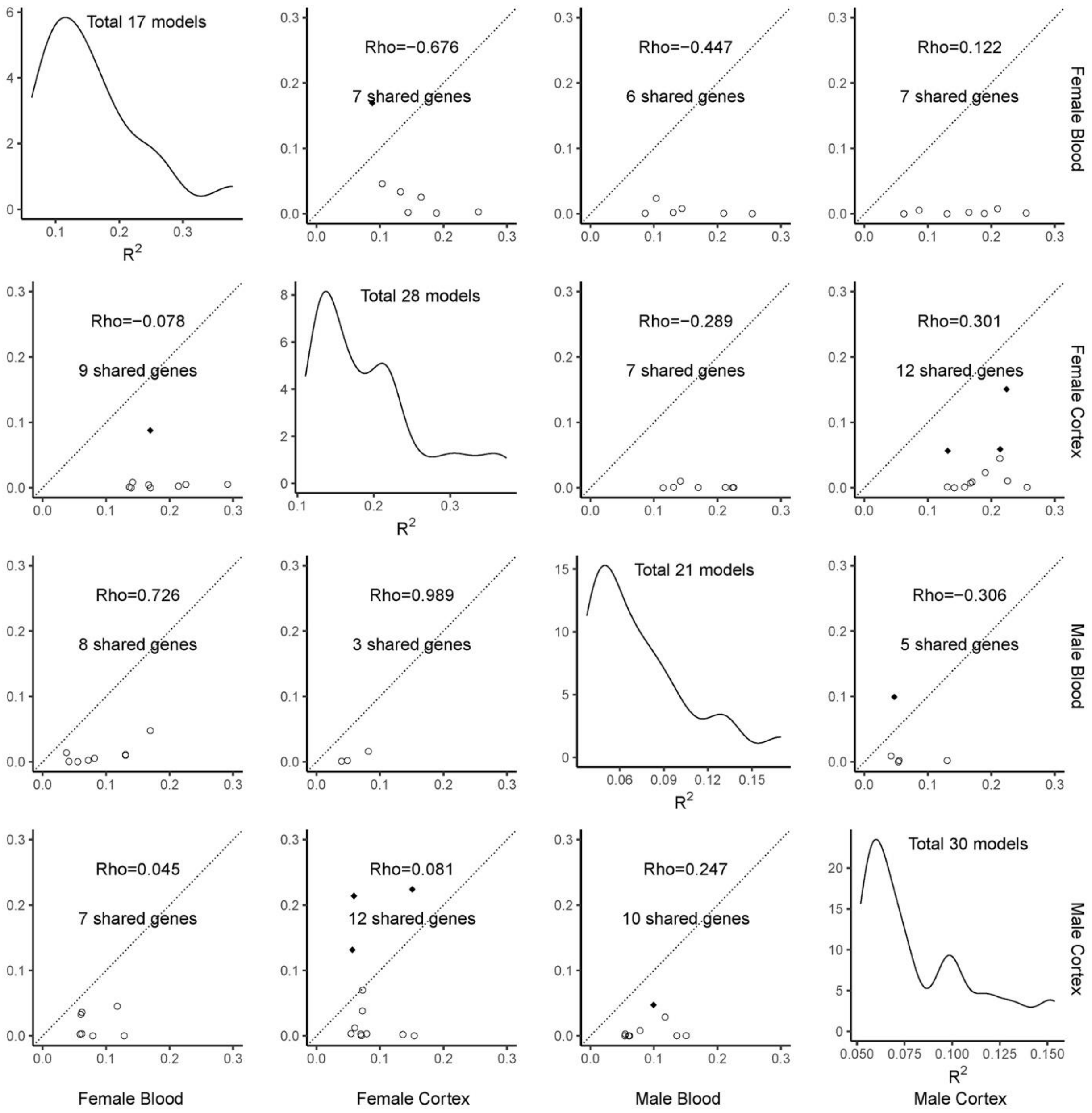


*Supplementary Figure 7. Comparison matrix of the balanced sex-stratified elastic net models trained in GTEx cortex and blood data, α=0.95. The number of genes being modeled in each tissue-sex set is listed in the distribution plot. The off-diagonal panels compared the $R^2$ of the models in pairs of tissue-sex sets. The four rows are for the female-blood, female-cortex, male-blood, and male-cortex. For each row, the genes of the significant models in the tissue-sex set were matched in the other sets for the matching $R^2$. The diamonds represent genes with a significant model ($R < 0.1$ and $p > 0.05$) in another tissue-sex set, while the circles represent genes with a non-significant model in another tissue-sex set. The number of gene models and the Pearson correlation coefficients were listed in the comparison plots.*

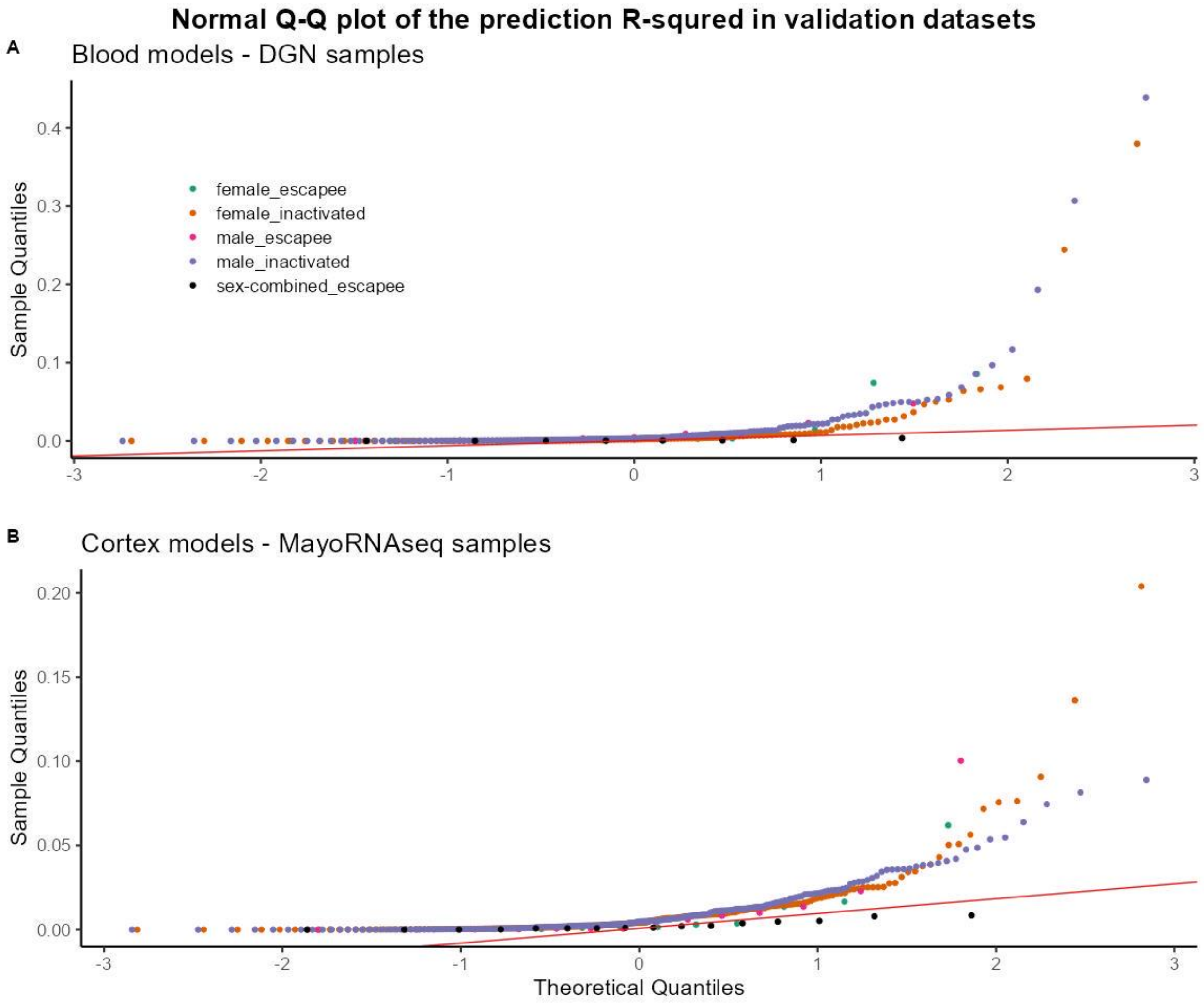


*Supplementary Figure 8. Q-Q plot of the prediction $R^2$ of blood (A) and cortex (B) models in validation datasets. All GTEx models were evaluated in these plots.*

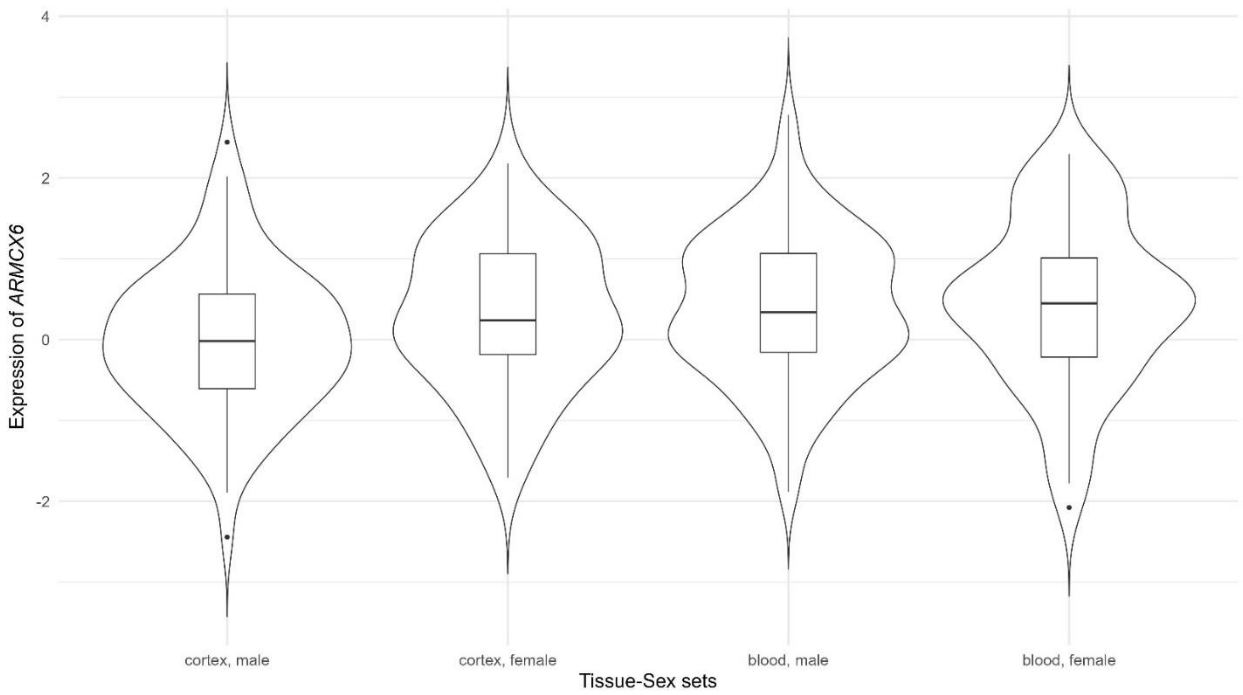


*Supplementary Figure 9. Distribution of the RNA expression level of the TWAS top hit, ARMCX6, in different tissue-sex sets*
